# Structural transition and antibody binding of Ebola GP and Zika E proteins from pre-fusion to fusion-initiation states

**DOI:** 10.1101/285130

**Authors:** A Lappala-Vernon, W Nishima, J C Miner, P W Fenimore, W Fischer, P T Hraber, M Zhang, B H McMahon, C-S Tung

## Abstract

Membrane fusion proteins are responsible for viral entry into host cells– a crucial first step in viral infection. These proteins undergo large conformational changes from pre-fusion to fusion initiation structures, and, despite differences in viral genomes and disease etiology, many fusion proteins are arranged as trimers. Structural information for both pre-fusion and fusion initiation states is critical for understanding virus neutralization by the host immune system. In the case of Ebola glycoprotein (GP) and Zika envelope protein (Zika E), pre-fusion state structures have been identified experimentally, but only partial structures of fusion initiation states have been described. While the fusion initiation structure is in an energetically unfavorable state that is difficult to solve experimentally, the existing structural information combined with computational approaches enabled the modeling of fusion initiation state structures of both proteins. These structural models provide an improved understanding of four different neutralizing antibodies in the prevention of viral host entry.

## Introduction

Enveloped viruses employ a common mechanism to enter the host cell (1). The first steps, receptor binding and membrane fusion, are initiated by the envelope protein (2–4). While specific details vary among different viruses, the envelope proteins invariably go through a large conformational change (5, 6) before initiating membrane fusion. These large conformational changes allow the envelope protein to assume an extended fusion initiation conformation: the envelope protein in the fusion initiation state is able to bridge across the viral- and the host membranes, subsequently bringing the two membranes in close proximity and starting the fusion process (7, 8). Viral neutralization by antibodies may involve binding to the fusion-state structure or inhibiting its formation. Therefore, viral envelope proteins are important foci for development of vaccines and therapeutics. Recent intense research focus on Ebola and Zika viruses has provided new data for structural modeling of these transitions.

Structural data for a number of viral envelope proteins are available in the Protein Data Bank (PDB) (9). Many of these known structures correspond to envelope proteins in the pre-fusion state, and some of the fusion-state structures only correspond to a partial molecule (usually in a low pH environment). To date, there are no structures for complete viral envelope proteins in the fusion initiation state; understanding the mechanics of the conformational change from pre-fusion to the fusion initiation state requires such a description.

Directly determining fusion-state structures for complete viral envelope proteins by experimental methods is difficult; molecular modeling offers a readily applicable alternative means to structural characterization. We describe the use of a knowledge-based methodology (homology modeling) to develop structures of viral envelope proteins in the fusion initiation state. We further extend the basic idea of homology modeling to include a simple concept “proteins and protein domains that fold similarly interact similarly”, as a result, developing structural models of envelope protein-antibody complexes. In this work we focus on envelope proteins from Ebola and Zika viruses. Ebolavirus causes Ebola hemorrhagic fever, a severe and highly lethal infection: the 2013-2015 West African Ebolavirus epidemic (December 2013 – 2015) resulted in approximately 11,000 confirmed deaths and 28,000 suspected cases (10). Zika virus, in contrast, causes a brief, relatively mild illness, but has been linked to congenital microcephaly and Guillan-Barré Syndrome in humans (11, 12); in mouse models, Zika virus causes microcephaly (13), as well as damage to the male reproductive system (14) and to adult neural stem cells (15). At the time of publication, eighty-four countries, territories and subnational areas report Zika transmission (16). Ebola and Zika viruses represent persistent threats to public health; there are limited options available for treatment or prevention of either virus. Although they are phylogenetically distinct viruses, their fusion subunits both adopt a trimer structure: in this study, we investigate some of the similarities, differences, and consequences of these fusion-state structures. We present the following models for Ebola GP and Zika E proteins:

1. A trimer model of Ebola GP in fusion initiation state with NPC1 receptor and neutralizing antibodies.
2. A trimer model of Zika E in the fusion initiation state with neutralizing antibodies and the surrounding 9-mer structure of Zika E protein in the pre-fusion state with neutralizing antibodies.

Our modeling approach is general and comprehensive, and can be used for developing structures of other pathogen proteins in their functional states for understanding their functions; the developed structure-based knowledge can further add to sequence-based information and improve vaccine design for viruses and other pathogens. Importantly, all of the developed models are testable experimentally, potentially leading to the discovery new targets for drugs and vaccines, and optimization of known vaccine- and drug targets.

### Ebola Envelope Protein GP1/GP2 and Zika Envelope Protein E

Ebola has a small (∼18-19 kb) negative-stranded RNA genome (17) that encodes eight viral proteins. Envelope glycoprotein (GP) is the viral surface protein responsible for host cell entry (18, 19) and has a sequence of 676 residues in all known Ebola strains. The N-terminus (residues 1-32) forms a signal peptide and the remaining protein residues (33-676) are collectively referred to as the GP portion. The host endoprotease furin cleaves GP into two segments: GP1 (residues 33-501) and GP2 (residues 502-676) (20). GP1 is responsible for receptor binding to the host cell (21, 22) while GP2 acts as a class I viral fusion protein (23). Several functional domains of Ebola GP are denoted in **Fig.** 1. Like Influenza and HIV envelope proteins, Ebola GP is a homotrimer in its functional state (24).

**Figure 1:**
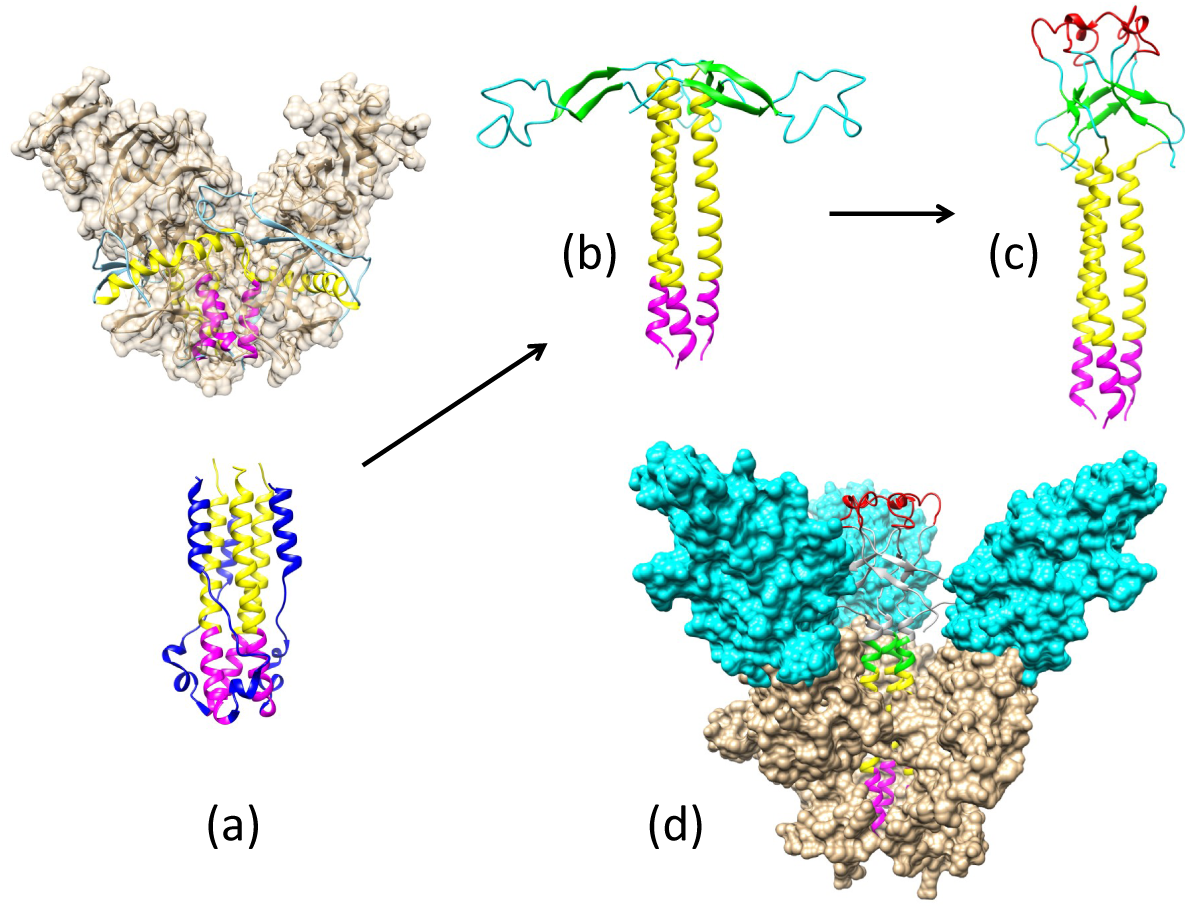
Structures of Ebola GP corresponding to the pre-fusion and fusion initiation states. **a.** (*Top*) A trimer structure of GP1/GP2 in the pre-fusion state and (*Bottom*) GP2 in the fusion initiation state were solved using x-ray crystallography (PDB accession codes 3CSY and 2EBO, respectively). **b.** Model of the pre-fusion structure of the GP2 trimer with a spring-loaded mechanism. **c.** Hairpin loop structure (residues 510 to 559) of GP2, including the fusion peptide (residues 524-539, *red*), remodeled into a conformation with the fusion peptide pointing toward the host cell surface. **d.** GP1/GP2 trimer in the fusion initiation state bound to the NPC1 receptor. Viral membrane is oriented at the bottom of each panel.

Since the 2015 epidemic, significantly more Ebolavirus sequence information has become available, including sequences of GP (more than 1,000 unique Ebola GP sequences can be found in the current NCBI Ebolavirus Database) (25). Significant efforts have been devoted towards solving the GP structure using various experimental approaches including x-ray crystallography (24, 26–28), NMR (29, 30) and Electron Microscopy (31, 32). Through this work, domain structures have been solved in different functional states, as has the trimer structure of the Ebola GP mucin-like region deletion mutant (GPmuc) in the pre-fusion state (24). All of these pieces of information contribute to a basis for developing a structural model of the GP in the fusion initiation state.

Zika virus is a member of the virus family Flaviviridae (33) that includes Dengue and West Nile viruses; it possesses a non-segmented, single-stranded, positive-sense RNA genome (10 kb) (34). The urgency in finding ways to combat the virus has increased significantly since a causal relation between Zika and apoptosis of human neurons was discovered (11). For EBOV, the envelope protein (GP) forms homotrimers that are distributed sparsely on the surface of the virus, as in influenza and HIV. By contrast, Zika Es form homodimers and cover the entire viral surface (the capsid comprises 90 homodimers, arranged as 30 rhombic faces with 3 dimers each) (35, 36). However, the partial structure of the Zika E in the fusion initiation state (at low pH) shows a trimer arrangement. Determining how Zika E transforms from a pre-fusion dimer to a fusion-state trimer presents a considerable challenge to the structural modeling community.

## Methods and Materials

### Homology modeling using a Motif-Matching Fragment Assembly method

Motif-Matching Fragment Assembly Method (MMFA) was used for complex homology modeling (described in our earlier work (37)). This approach was used to develop structural models of neutralizing antibodies binding to its targeted fusion proteins in their functional unit and state.

### Structure alignment and superposition

To develop fusion initiation state structure of the GP protein using the spring-loaded mechanism, domain structural assembly was accomplished by superposition of structurally matched regions (e.g., helical segments). The MatchMaking option in Structural Comparison Tool of Chimera (38) is used for structure alignment and superposition of the structurally matched segments.

### Structural refinement

Molecular dynamics simulations of Zika and Ebola models were performed in a standard manner as described in Hess et al. (39). Initially, structures were energy-minimized with a steepest descent minimization algorithm, after which the protein underwent a 100 ps NVT (constant number of particles, volume and temperature) equilibration step, followed by NPT (constant number of particles, pressure and temperature) equilibration for another 100 ps. The production run was performed under standard NPT conditions at 300K for 100 ns.

### Graphics

Molecular graphics images were produced using the UCSF Chimera package (40) from the Resource for Bio-computing, Visualization and Informatics at the University of California, San Francisco.

## Results

### Pre-fusion and fusion initiation state structures of EBOV GP (‘spring-loaded model’)

The crystal structure of EBOV GP1/GP2 has been described in the pre-fusion state (accession code: 3CSY) (24). In this structure, GP1 and GP2 exist as a trimer, with the GP2 bound from the outside and the inside of the GP1 proteins. A partial structure of GP2 (lacking N-terminal and C-terminal domains) in the fusion initiation state has also been solved at low pH (accession code: 2EBO) (26). The pre-fusion and fusion initiation states of GP2 show different folds (see **Fig.** 1a) with a small triple-helix structure shown in both (highlighted in magenta).

In the pre-fusion structure, the yellow/magenta regions of GP2 are wrapped around the GP1 trimer on the outside of the GP1 trimer. For the yellow helices to stack on top of the magenta helices as indicated (PDB: 2EBO), they must rearrange from the outside to the inside of the GP1 trimer. A close inspection of the GP structure in the pre-fusion state reveals a connection between two GP1 proteins in the trimer through loops S90 to P93 and P126 to R130. To accomplish the transition step, this weak bridge needs to be disconnected to allow the N-terminal part of GP2 to move from the outside of the GP1 trimer to the inside.

The C-terminal region of GP2 (highlighted in blue, **Fig.** 1a) needs to ‘peel off’ from the central triple helix of the fusion initiation state in order for GP2 to bind and fit into the internal part of the GP1 trimer. Since the C-terminal end of GP2 contains the trans-membrane domain that points toward the viral surface – and is less likely to be immunogenic – our model is truncated at residue 599. To account for the large conformational changes during the pre-fusion to fusion transition, a spring-loaded model is applied (41, 42). This same model was considered as the mechanism for the Ebola GP structural transition in a previous study by White et al. (43), and is supported by structural information of Ebola GP in two different states (**Fig.** 1a).

The spring-loaded model suggests that the two GP2 helices in the pre-fusion state structure (3CSY) should be stacked together to form one long helix, as shown in the fusion initiation state structure (2EBO). When the two helices become one long helix, the N-terminal region of GP2 (residues 502-555) will be stacked on top of the long triple helix and pointed radially outward (**Fig.** 1b).

The known structures of viral envelope proteins in the fusion initiation state usually show a trimer arrangement in a compact state (6) where the fusion peptides are tightly localized on top of the trimer and exposed to the host cell membrane (44). The GP2 structure (**Fig.** 1c) has the fusion peptides (residues 514-539, highlighted in red) separated and aligned outwards, parallel to the viral membrane, instead of upward toward the host cell membrane. In order to develop a model that matches known fusion initiation state structures, the sole *β*-hairpin structure in GP2 (residues 517-520, 543-546) are fitted to a trefoil fold *β*-hairpin template (PDB: 2F2F) at each trimer. The modeled *β*-hairpins in the trefoil fold are then docked on top of the yellow triple helix using the disulfide bond between C-511 and C-556 as a constraint. Finally, the low pH fusion loop structure (PDB: 2RLJ) is used as a template to model the fusion peptide structure (residues 521-542). The final structure of the GP2 trimer in the fusion initiation state is shown in **Fig.** 1c.

Niemann-Pick C1 (NPC1) has been identified as the entry receptor for EBOV GP (45), which must be primed to a fusion-competent state before binding an NPC1 receptor. The priming process includes cleavage of the mucin-like domain, the glycan cap and the outmost strand of the proposed receptor binding region (24, 46). Using the NPC1-bound GP1 structure (PDB: 5F1B) as a template, primed GP1-GP2-NPC1 complex in the fusion initiation state is modeled (**Fig.** 1d).

Based on these results we propose that the connecting bridge in the trimer (S90 to P93, P126 to R130) serves as a gate for the pre-fusion to fusion structural transition. If this gate is locked, the structural transition step is inhibited, which impedes viral replication.

### Pre-fusion and fusion structure of the Zika E

The basic unit of the Zika E protein, based on structures derived from x-ray crystallography and cryo-EM (PDB accession codes: 5JHM, 5LBS, 3CSY), shows the protein in a head-to-tail dimer arrangement (**Fig.** 2a). While the structure of Zika E in the fusion initiation state is not known, the partial Zika E structure of the related Dengue virus (DENV), has been described in its fusion initiation state (PDB accession code: 1OK8). Due to the high sequence similarity of DENV and Zika, it is a simple exercise to model the structure of Zika E in the fusion initiation state using the homology modeling approach (**Fig.** 2b).

**Figure 2:**
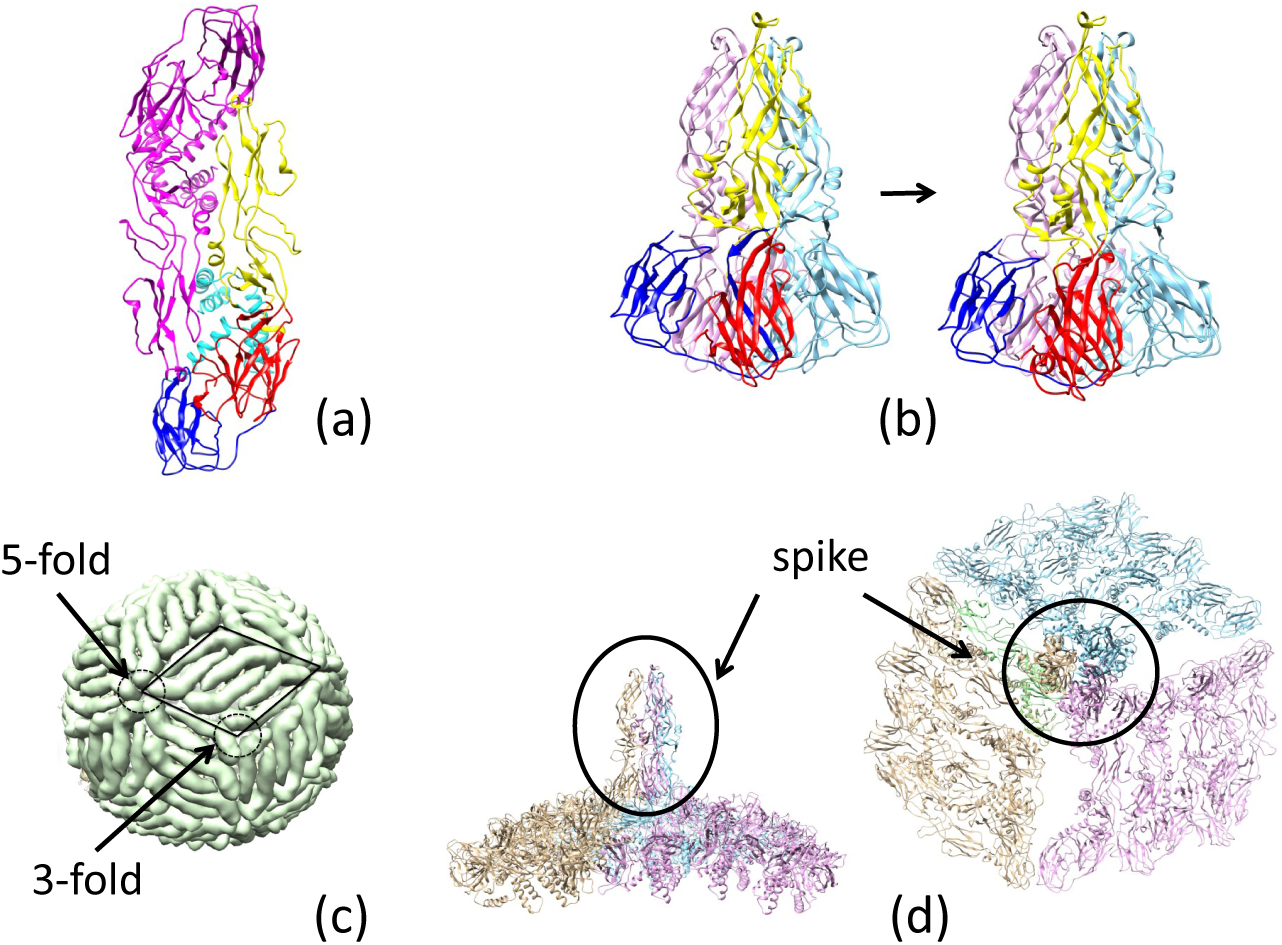
The pre-fusion and fusion structures of the Zika E protein. **a.** The 3.8 Å ngström cryo-EM structure of the Zika E dimer. **b.** *Left* Partial structure of DENV E, and *Right* modeled structure of Zika E in their fusion initiation states. These structures are divided into four domains, highlighted as follows: (I: red, II: yellow, III-1: blue, III-2: cyan). **c.** The 180 copies of Zika E cover (from cryo EM) the surface of the viral capsid, with intersection points of 3- and 5-fold symmetries marked. **d.** A model of side- and top-views of the Zika E trimer (marked as a spike) in the fusion initiation state.

Based on the known structure, Zika E can be divided into three domains: domain I (residues 1-52, 132-193, 280-296), domain II (residues 53-131, 193-276) and domain III (residues: 297-501). In this work, we further divide domain III into domain III-1 (residues 297-403) and domain III-2 (residues 404-501). Domain III-2 is absent in the fusion initiation state structure. Converting Zika E from 2-fold to 3-fold symmetry is a significant modeling challenge. Schematic representations of the flavivirus membrane-fusion mechanism have been proposed (47, 48), involving (i) a low-pH induced dissociation of the Zika E dimer, (ii) outward projection of Zika E monomer, and finally, (iii) formation of the fusion-state trimer. The proposed process does not provide information about the full Zika E structure in the fusion initiation state, so the relative position and orientation of the trimer with respect to the rest of the Zika E monomers on the surface of the virus requires further elucidation. Here, we propose a direct path for the Zika E pre-fusion to fusion structural conversion.

The cryo-EM structure of Zika E (PDB: 5IRE) is shown in **Fig.** 2c. In this representation, three dimers form a rhombus-shaped hexamer (highlighted by the parallelogram), and thirty copies of this hexamer are sufficient to cover the entire surface of the virus. These hexamers form 3- and 5-fold symmetries at their vertices, where the ends of either three or five Zika E proteins meet. The basic unit used to model the pre-fusion to fusion transition are three dimers that form a 3-fold symmetry. Since domain III-2 is absent in the DENV E protein fusion initiation state structure, we propose that this region serves as an anchor for the structural transition, and domains I, II and III-1 from each of the three Zika E molecules are involved in the structural transition. This transition is accomplished through a 90° rotation of domain I/II. The binding modes of I/II and III-1 are different between the pre-fusion (3CSY) and fusion (1OK8-derived) structures. The I/II and III-1 binding mode for 1OK8 upsets the connection between III-1 and III-2, thus the I/II/III-1 arrangement of the pre-fusion structure is used in the final fusion initiation state (spike in **Fig.** 2d).

### Neutralizing antibody blocks viral entry by pre-fusion and fusion initiation state interactions

Neutralizing antibodies inhibit pathogen entry into the host cell. Due to the fact that this is a complex process (49), blocking viral host-cell entry may occur at multiple stages. Since the conformational transition of the viral envelope protein from a pre-fusion to a fusion initiation state is a crucial step in host-cell entry, we explore different neutralizing antibodies, and their roles in blocking cell entry, by studying the binding of antibodies to Zika E in pre-fusion and fusion initiation states.

### Antibody KZ52 blocks EBOV entry by preventing GP structural transition from a pre-fusion to a fusion initiation state

The KZ52-bound Zaire-Ebola GP trimer in the pre-fusion state has been solved using x-ray crystallography (PDB accession code: 3CSY). Residues of the GP that are responsible for KZ52 binding include GP1(residues 42/43) and GP2(residues 505-511/513-514/549-553/556). If the N-terminal portion of GP2 cannot peel off from the outside and move to the inside of the GP1 trimer then the conformational transition to the fusion initiation state structure cannot be completed. Therefore, binding of KZ52 to EBOV GP prevents the structural transition of the protein to the fusion initiation state. A similar binding mechanism was observed for the binding of 16F6 to Sudan-Ebola GP (PDB accession code: 3S88). Therefore, we may argue that 16F6 blocks cell entry by preventing the structural transition of GP from a pre-fusion to a fusion initiation state.

### Antibody mAb100 blocks EBOV entry through two different mechanisms

Antibody mAb100, when used in conjunction with mAb114, can protect non-human primates against all signs of Ebola virus disease (50). The crystal structure of mAb100-bound EBOV GP (PDB accession code: 5FHC) shows that mAb100 binding occludes access to the cathepsin-cleavage loop **Fig.** 3, thus preventing viral entry. It is a straightforward exercise to develop a model of Ebola GP trimer with bound mAb100 by superimposing the GP structure in 5FHC with that in 3CSY. In this model, mAb100 interacts with two copies of GP2 in the trimer. As a result, mAb100 directly blocks the transition from pre-fusion to fusion initiation state. From the perspective of 5FHC, mAb100 recognises and binds to a loop near the fusion peptide in GP2. Using this binding mode, we develop a model of mAb100 binding to a GP trimer in the fusion initiation state. It was further shown that up to three mAb100 can bind simultaneously to a GP trimer in the fusion initiation state. While the fusion peptide is exposed when the GP trimer is in the fusion initiation state structure, the exposure is blocked when mAb100 binds to the viral GP.

**Figure 3:**
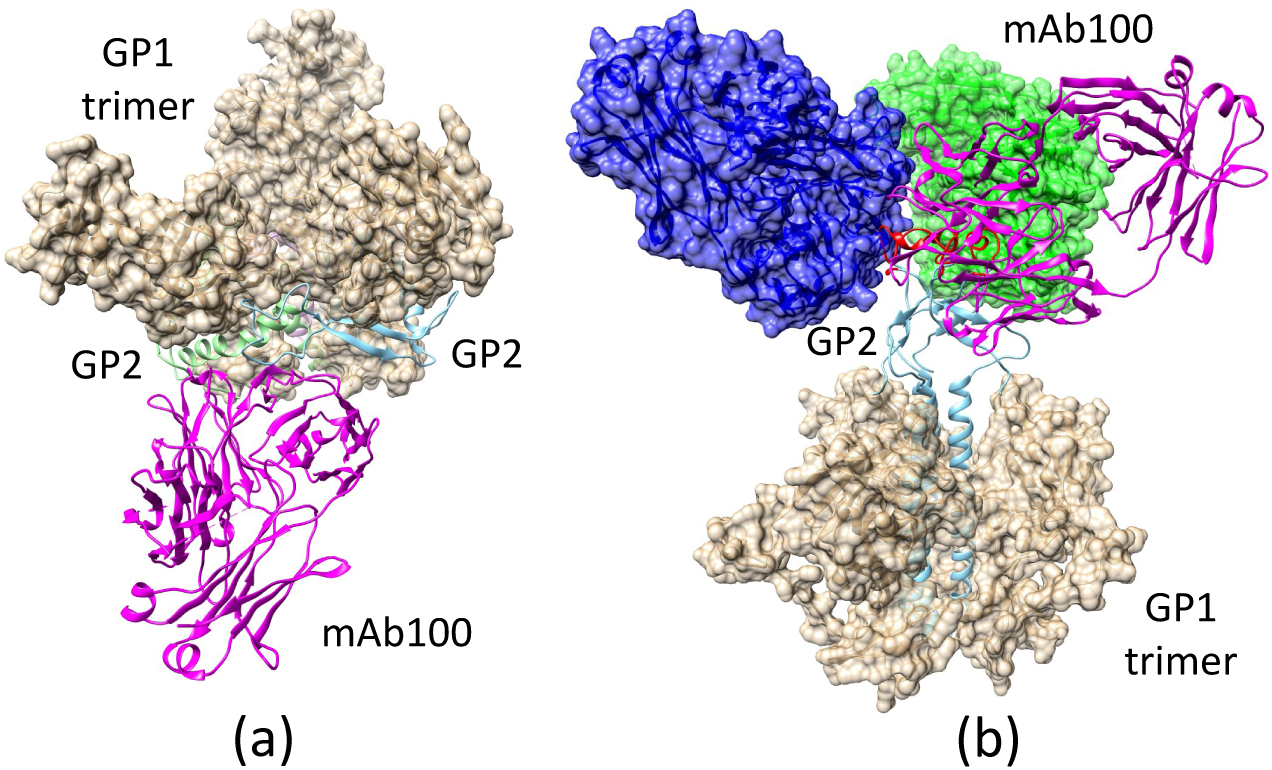
The mAb100 antibody can bind to the EBOV GP trimer in both pre-fusion and fusion initiation states. **a.** Modeled mAb100 binding to two GP2 monomers when the GP trimer is in the pre-fusion state, preventing the fusion initiation state transition. **b.** A post-transition model, where the mAb100 binding site on GP2 is exposed at the top of the trimer, allowing the antibody to bind. The model shows that up to three mAb100 antibodies can bind to the GP trimer in the fusion initiation state structure without steric interference.

### Antibody 2A0G6 binds to Zika E in the fusion initiation state and blocks the exposure of the fusion peptide

The structure of Zika E, bound with a broadly-protective antibody 2A0G6 (**Fig.** 3b) has been solved using x-ray crystallography (PDB accession code: 5JHL) (51). The structure shows that the antibody interacts with Zika E residues 76, 77 and 105-108. These residues lie on the tip of the GP close to the fusion loop. In the pre-fusion state, these residues are buried and cannot be reached by the antibody, but in the fusion initiation state, these residues become exposed. When the antibody binds, the fusion loop (residues 100-108) is blocked from exposure, and the fusion process is blocked.

### Antibody EDE1-C8 blocks the pre-fusion to fusion transition of Zika E and prevents cell entry

The antibody (EDE1-C8) binds to the pre-fusion state structure of Zika E on the surface of the virus as shown in **Fig.** 4. With the antibody bound, it prevents the peeling off of the N-terminal regions (I, II and III-1) of the Zika E protein from its neighboring Zika E and forming the fusion-state trimer structure. With the Zika E locked in the pre-fusion structure, it prevents the pre-fusion to fusion transition.

**Figure 4:**
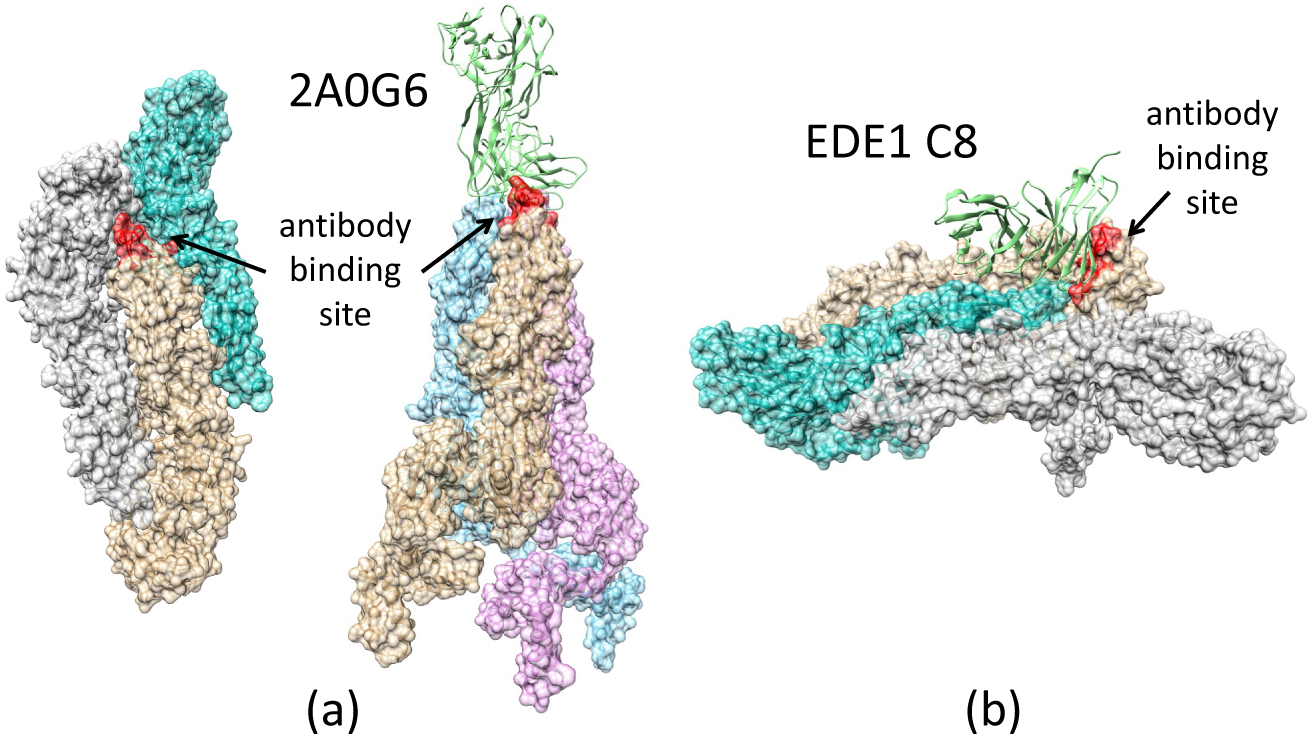
Two different antibodies, 2A0G6 and EDE1 C8, interact with the Zika E to prevent viral cell entry with different mechanisms. In this figure, the antibody recognition site is colored in red. **a.** Zika E in the pre-fusion state, the antibody-binding site is partially buried and prevents binding of 2A0G6 to E. **b.** Zika E in the fusion initiation state, antibody-binding site is exposed, allowing 2A0G6 to bind to the E protein. The antibody EDE1 C8 binds to two copies of the Zika E and prevents transition from the pre-fusion to fusion initiation state.

### Videos describing the transitions from the pre-fusion to fusion structures

Large conformational changes are associated with the pre-fusion to fusion initiation state structures of the viral envelope protein. Even though the structures at both ends of the transition are known, it is not obvious how to transform the structures from one to the other. To illustrate the complex process, two videos as an artists representation of the transition were developed. Each of the two videos depicts a possible (but not exclusive) transition pathway for Ebola GP and Zika E respectively. In these videos, the integrity of stereochemistry was maintained for structures in each frame. Many of the features associated with the transition and described in the manuscript are easily identifiable in the videos.

## Conclusions

Viral cell entry is a complex process involving multiple molecular participants and conformational changes. We provide a plausible mechanism in atomic detail for the pre-fusion/fusion-state transition for both Ebola and Zika viruses. While these two viruses are very different, their fusion proteins share common features in the initiation of cell entry. The most obvious common feature is a trimer arrangement of the proteins in the fusion initiation state. This can be seen in other viruses, such as Influenza and HIV. Secondly, these fusion proteins undergo large conformational changes to transition from pre-fusion to the fusion initiation state structure. To prevent viral cell entry, antibody can act on the fusion protein using two different mechanisms: 1) preventing the structural transition, and 2) blocking fusion peptide access to the cell membrane With the known structures of these proteins in the pre-fusion state and partial structures of the proteins in the fusion initiation state, we developed near-complete fusion-state structural models of Ebola GP and Zika E. Using a spring-loaded mechanism proposed for influenza hemagglutinin (HA) structural transition, the fusion initiation state trimer structure of the Ebola GP was developed. The fusion initiation state trimer structure of the Zika E was developed from three dimers on the surface of the virus utilizing the three-fold symmetry.

Combining the information of Ebola GP/Zika E structures in both the pre-fusion and the fusion initiation states as well as the structures of these proteins in complex with different neutralizing antibodies, we investigated the mechanisms used by the antibody to prevent the process of viral cell entry. We found that KZ52 (Ebola) and EDE1-8 (Zika) prevented cell entry by blocking the pre-fusion to fusion structural transition. We proposed that the connecting bridge in the GP1 trimer (S90 to P93, P126 to R130) serves as a gate for the pre-fusion to fusion structural transition: if this gate is locked, the structural transition step is inhibited and the viral life cycle is halted. We also demonstrated that the antibody 2A0G6 binds to the Zika E in the fusion initiation state and prevents the fusion peptide from reaching the cell membrane. The antibody mAb100 is able to bind Ebola GP in both the pre-fusion- and the fusion initiation states, preventing the viral cell entry both by inhibiting structural transition and by blocking fusion peptide access to the cell membrane.

With the collective efforts to expand the information base in both biological sequences and structures, GenBank and PDB provide invaluable information. This information enables the expansion of structural modeling to a new realm that includes proteins in higher-order structure and/or transition. Using Ebola GP and Zika E as examples, we have demonstrated that a conceptually straightforward, knowledge-based approach allows us to develop models of the fusion initiation state structures for both proteins in their functional units. With this information, we were able to study the binding of neutralizing antibodies to these proteins and proposed the exact mechanisms for how these antibodies block cell entry to stop viral replication.

## Author contributions

All authors were directly involved in regular discussions of model development and provided direction throughout the project as well as participated in manuscript writing and editing. AL-V provided the energy-optimized structures of all the models developed from this work. She is also responsible for combining different edited versions into the final manuscript. WN developed the two videos that demonstrated the transition of the two proteins from pre-fusion to fusion state. C-ST did the original structural modeling work and provided oversight of the whole project.

## Acknowledgments

We gratefully acknowledge the support of the U.S. Department of Energy through the LANL/LDRD Program, project number 20170509DR for this work. C-S T. is a LANL retired scientist and worked on this project on *a pro bono* basis. M.Z. acknowledges the funding of NIAID Centers of Excellence for Influenza Research and Surveillance (CEIRS), contract number HHSN272201400004C.

## SUPPLEMENTARY MATERIAL

### Modeling fusion state structure of the Ebola GP using the spring-loaded mechanism

The crystal structures of the Ebola GP in fusion and pre-fusion states are shown as the top and the bottom images in **Fig. S1** respectively. Neither of the structures is complete: the pre-fusion GP structure (3CSY) is significantly more complete and includes large portions of GP1 (residues 31 to 310) and GP2 (residues 502-595). Only partial structure of GP2 (residues 557 to 630) is solved and available to the public (2EBO). The color coding of the helical regions corresponds to the spring-loaded mechanism with the following color assignments: green, 560-564; yellow, 565-582; magenta, 583-595.

In the fusion state structure of the GP trimer, the C-terminal regions (residues 596 to 603, highlighted in red) are wrapped around the triple-helices formed from the N-terminal regions (residues 557-595) of the three GP2 molecules. In the pre-fusion structure of the GP trimer, only a small portion of GP2, the triple-helix formed from the magenta-colored helices, is internal to the GP1 trimer (shown in a space-filling model in **Fig. S1a**). The remaining parts of GP2 are wrapped around the GP1 trimer from the outside. With the central cavity in the GP1 trimer (3CSY) being limited and only being able to accommodate a tight triple helix, the C-terminal regions of GP2 (red helix in **Fig. S1a**) has to be ‘peeled’ from the central triple-helix and move to the space below the triplex. With this part of the molecule including the membrane proximal external region (MPER) and the transmembrane helix (TM), its main function should be anchoring the protein to the surface of the virus. With insufficient information to model the structure of this portion of the molecule, we decide to leave this segment out of our model.

The common structure between the pre-fusion (3CSY) and fusion (2EBO) structures of GP2 is the triple helix formed by segments (residues 583-595) boxed and color highlighted in magenta as shown in **Fig. S1b**. By matching the magenta-colored helices, the structure of the three helical segments (magenta, yellow, green) in 3CYS can be replaced by a single long helix from 2EBO. The N-terminal residues (514 to 564 color-coded in cyan and green) of GP2 from 3CSY are added to the long helix by matching the green segments of the helices (see **Fig. S1c**) resulting in the model of GP2 in the fusion state (see **Fig. S1d**).

### Remodeling the fusion loop structure

With the modeled structure as shown in **Fig. S1d**, in a trimer arrangement, the fusion loops are pointing sideways instead of upwards to the host membrane. We decide to remodel this portion (residues 514 to 559) of GP2 in the fusion state structure. With the segment adopting a *β*-hairpin structure (see **Fig. S2a**), the task becomes arranging three *β*-hairpins into a trefoil fold. By analyzing the protein database, we have learned that the structure of a *β*-hairpin trefoil fold can be found in the crystal structure of the cytolethal distending toxin (CDT) (PDB accession code: 2F2F, see the top image of **Fig. S2a**). By matching the two *β*-strands in the GP2 hairpin to those in the trefoil fold, one can model the hairpin in a trefoil fold arrangement as shown in **Fig. S2b**. The NMR structure of the fusion loop (residues 525 to 538) has been solved and available in PDB (accession code: 2RLJ and shown as the red molecule in **Fig. S2c**). We substituted this known structure of the fusion loop into the corresponding region in the hairpin (colored in blue in **Fig. S2c**) and obtained the remodeled structure of the hairpin as shown in **Fig. S2d**. The final model of the GP2 trimer in the fusion state is shown in **Fig. S2e**.

**Figure S1:**
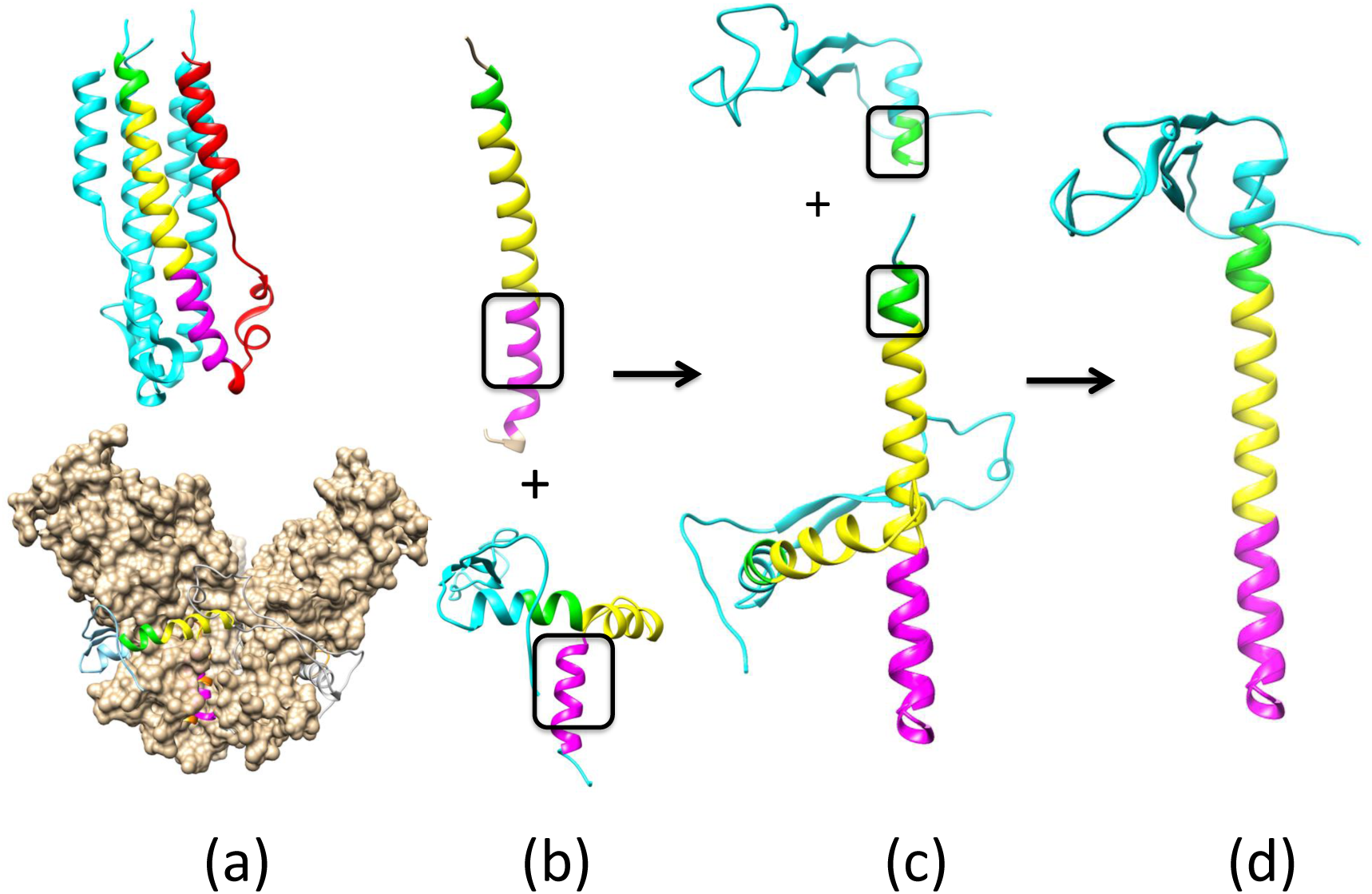
**a.** Incomplete crystal structures of the Ebola GP in its pre-fusion state (above) and the fusion state (below). **b.** Modeled triple helix formed by segments (residues 583-595) boxed and highlighted in magenta. **c.** N-terminal residues (514 to 564 color-coded in cyan and green) of GP2 from 3CSY added to the long helix by matching the green segments of the helices. **d.** The resulting complete model of the GP2 in the fusion state.

### Determination of Zika E domain III-1 placement in the fusion-state structure

It is obvious that domains I/II (residues 1 to 304, colored in cyan in **Fig. S3a** and **Fig. S3b**) need to rotate 90° to transit from the pre-fusion to the fusion state. It is also clear that domain III-2 (residues 404 to 501, colored in green in both **Fig. S3a** and **Fig. S3b**) should serve as the anchor for the Zika E in the fusion state and should be connected to other E proteins on the surface of the virus. Placing the domain III-1 (residues 305-403) relative to other domains in Zika E when the protein is in the fusion state is a problem that needed to be addressed: in the first case, when domains I/II move into the fusion state structure, domain III-1 can move together with domains I/II (as shown in **Fig. S3a**); in the second case, domain III-1 can stay with domain III-2 and not move together with domains I/II (as shown in **Fig. S3b**). In these figures, the connecting residues between domain II/III-1 and III-1/III-2 are highlighted as orange spheres. One can see that in case one (**Fig. S3a**), the distance between the two orange spheres is ∼50 Å, which is too long to form a connection. If domain III-1 did not move together with domains I/II as in the second case, the two orange spheres that connect domains II/III-1 and III-1/III-2 is separated with appropriate distances to form the connection. Based on this exercise, we determined that domain III-1 should stay with domain III-2 while domains I and II is rotated into a new position when Zika E protein undergoes the pre-fusion to the fusion state transition.

**Figure S2:**
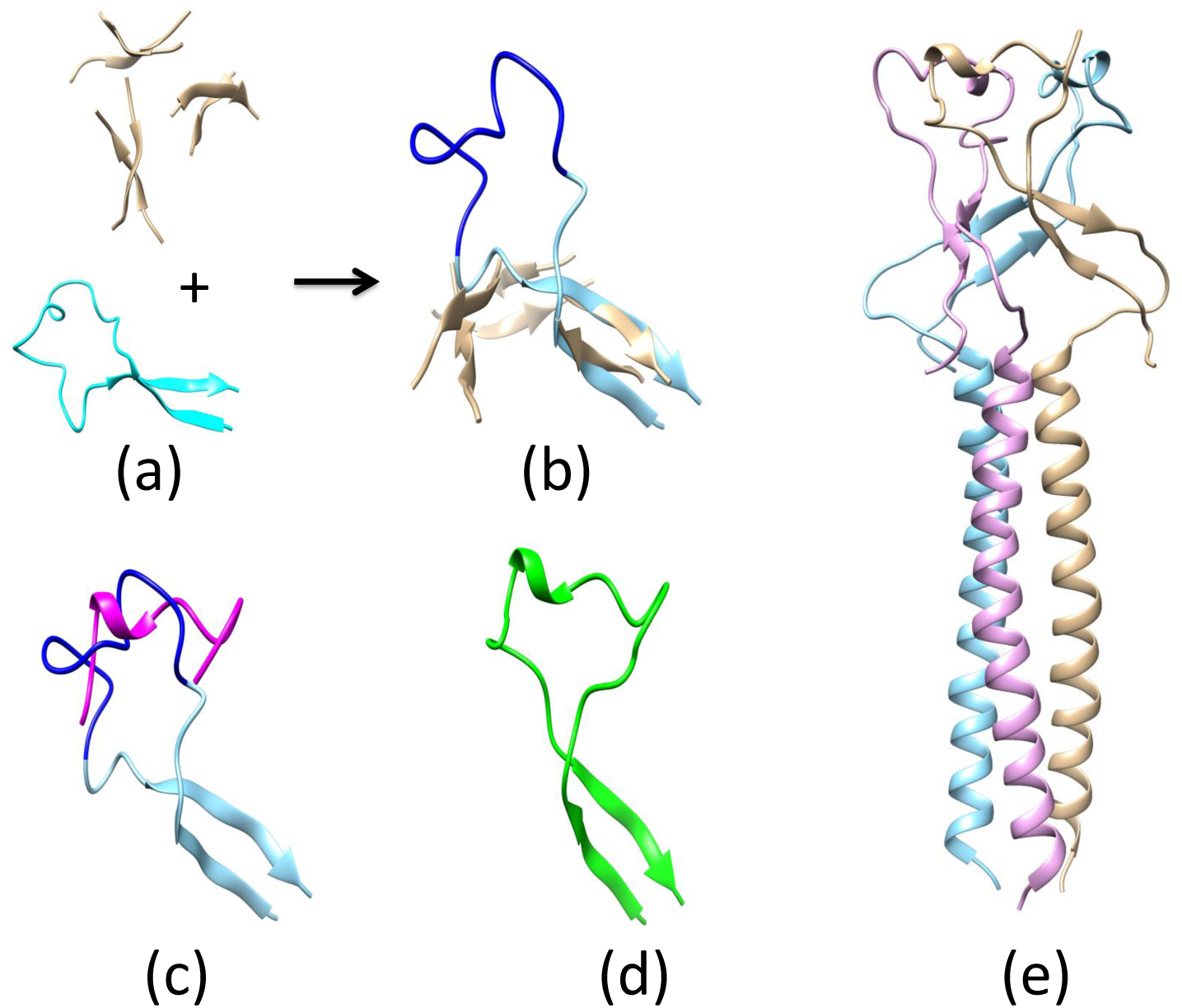
**a.** Residues 514-559 (below). Protein database analysis shows that the structure of a *β*-hairpin trefoil fold can be found in the crystal structure of the cytolethal distending toxin (CDT) (above) **b.** Re-modeled residues 514-559 of GP2 in the fusion state structure adopt a *β*-hairpin structure. These three *β*-hairpins form a trefoil fold. **c.** The NMR structure of the fusion loop (residues 525 to 538) shown in red. The known structure of the fusion loop was substituted into the corresponding region in the hairpin (shown in blue) **d.** The remodeled structure of the hairpin. **e.** The resulting complete model of the GP2 in the fusion state.

### Binding of EDE1 C8 to Zika E proteins on the surface of the virus

While the crystal and solution environments are quite different, the known structures of various proteins in the two different environments have shown similar structural folds (52). Here, we assume that the binding mode derived from crystal structures of the protein/antibody complexes provides a stable interaction for the complex under physiological conditions. The crystal structure of EDE1 C8/Zika E complex has been solved (PDB accession code: 5LBS) and is shown (ribbon model) in **Fig. S4a**. We can use this binding mode to tell how the antibody is going to interact with the Zika E when it is in its functional state (i. e when the E proteins cover the entire viral surface). In this figure, three Zika Es in the functional state (PDB accession code: 5IRE) are shown in a space-filling model (blue, cyan and gray respectively). The task is to bring the Zika E/antibody complex to the surface of the virus. When we match the Zika E in the Zika E/antibody complex (shown in **Fig. S4a**) to one of the Zika E monomers (colored in blue) on the surface of the virus, we can bring the antibody to the surface of the virus as shown in **Fig. S4b**. As a result of this exercise, we see that EDE1 C8 is interacting with two E proteins (colored in blue and cyan) as shown in **Fig. S4c**.

**Figure S3:**
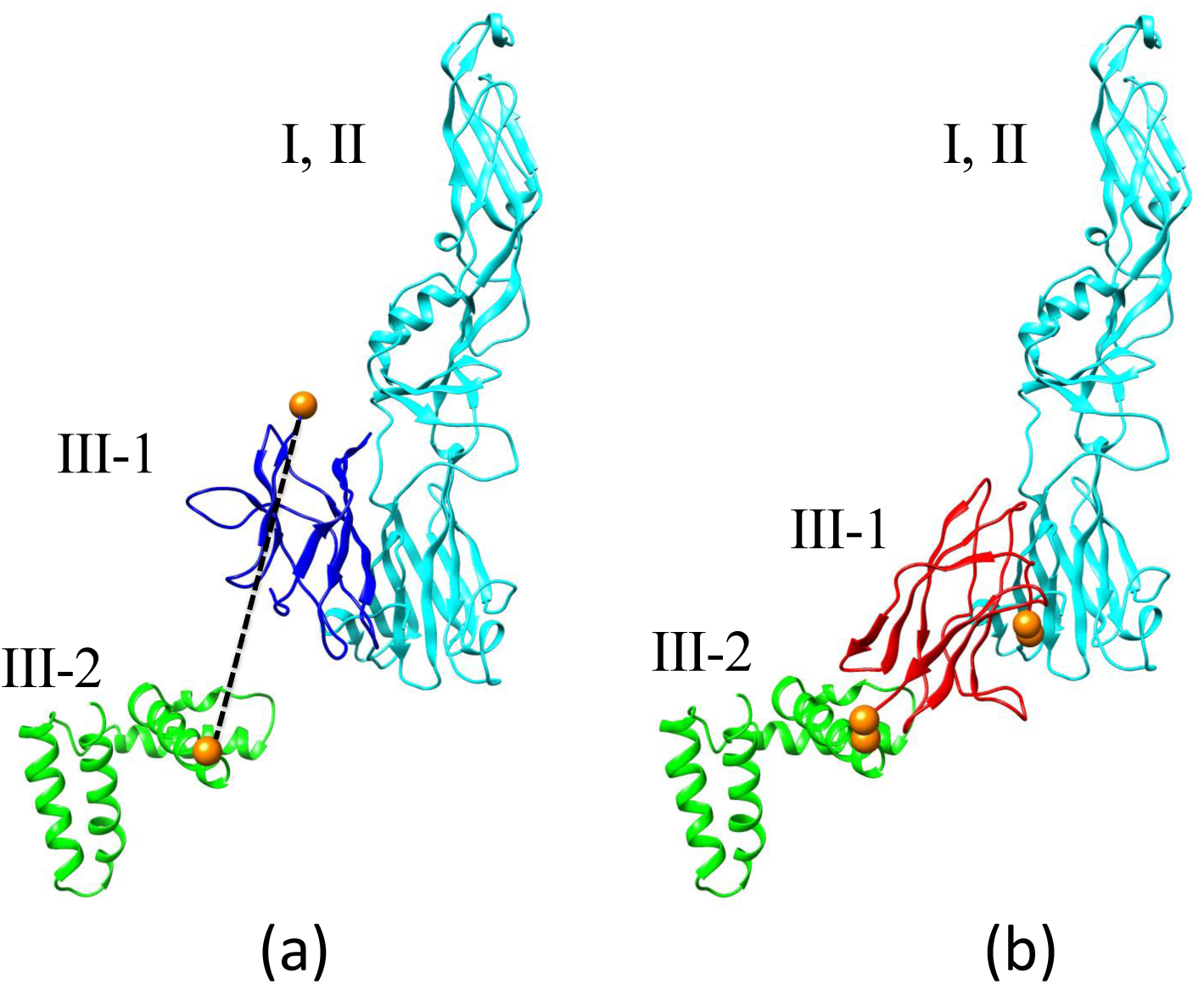
**a.** Zika E Domains I and II (residues 1 to 304) shown in cyan, domain III-1 (residues 305-403) shown in blue, domain III-2 (residues 404-501) shown in green. **b.** Zika E Domains I and II (residues 1 to 304) shown in cyan, domain III-1 (residues 305-403) shown in red, domain III-2 (residues 404-501) shown in green.

### Animation

**Video 1.** A transition path of Ebola GP between pre-fusion and fusion states (*left*: side, *right*: top). The intermediate structures were made by morphing and minimization. Each domain is colored as follows– GP1: yellow, red and blue, GP2: orange, cyan and green).

**Video 2.** A transition path of Zika E between pre-fusion and fusion states (*left*: side view, *right*: top view). To clarify the structure, Zika domains were colored as follows– I: yellow, II– red, III-1: blue, III-2: cyan, and the remaining two units of the trimer orange and green. the remaining structures are colored in gray.

**Figure S4:**
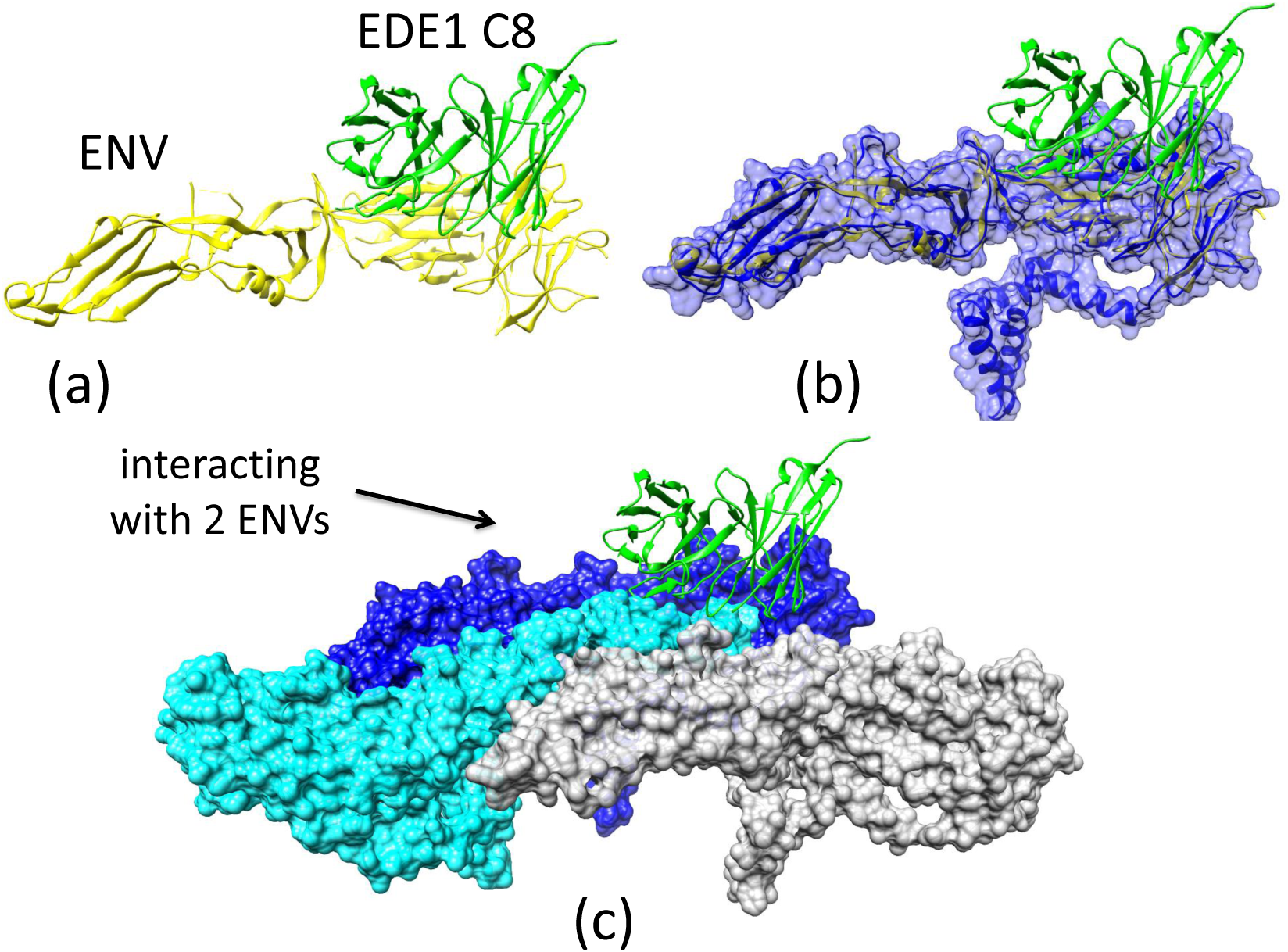
**a.** The crystal structure of EDE1 C8/Zika E complex (PDB accession code: 5LBS) is shown as a ribbon model. **b.** After matching Zika E in the virus/antibody complex to one of the Zika monomers on the surface of the virus, the antibody can be positioned as shown. **c.** EDE1 C8 interacting with two Zika Es (colored in blue and cyan).

