## Supplementary figures and images for "Structural transition and antibody binding of Ebola GP and Zika E proteins from pre-fusion to fusion-initiation states"

### Supplementary Materials

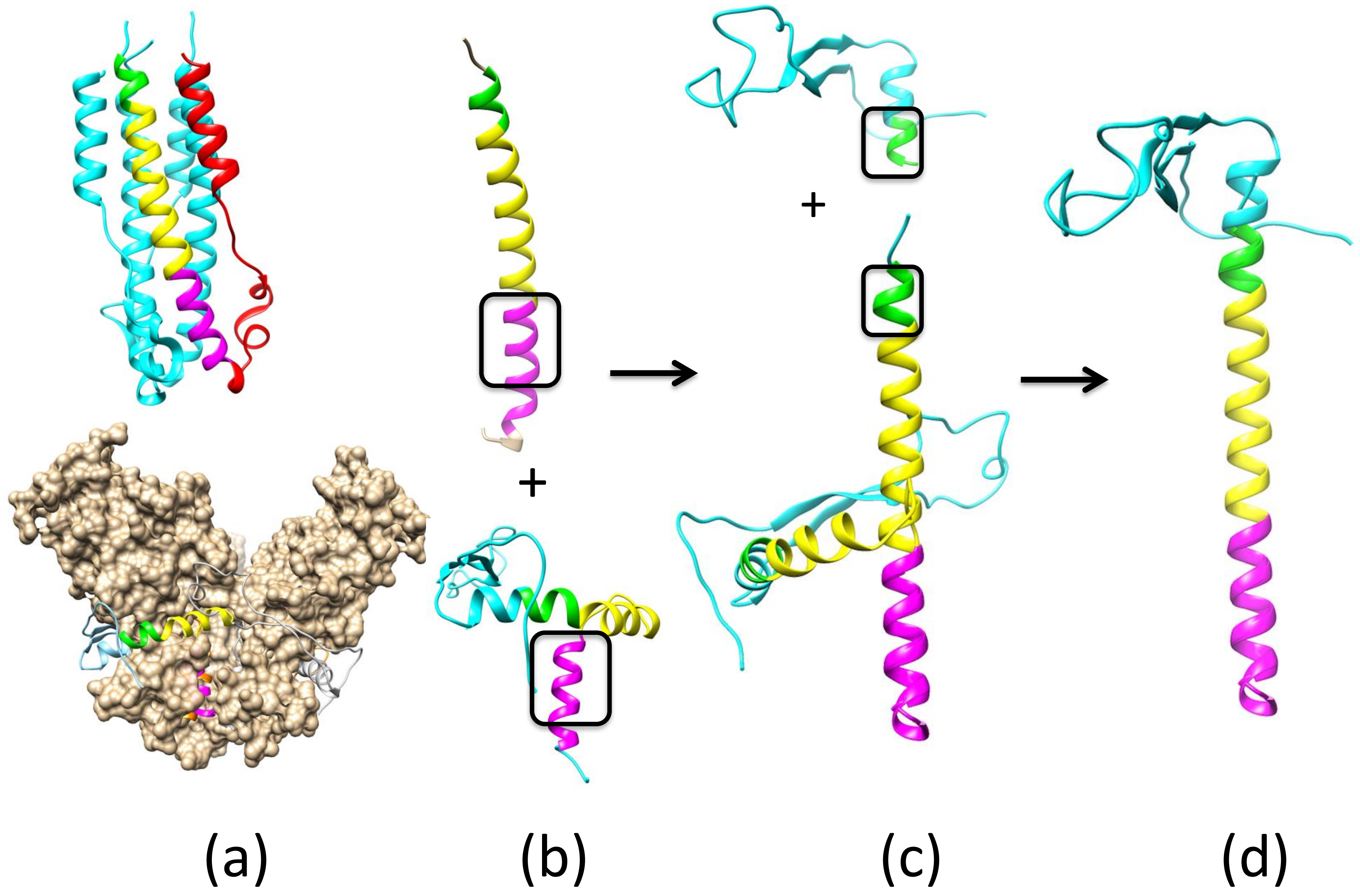

### Supplementary Materials

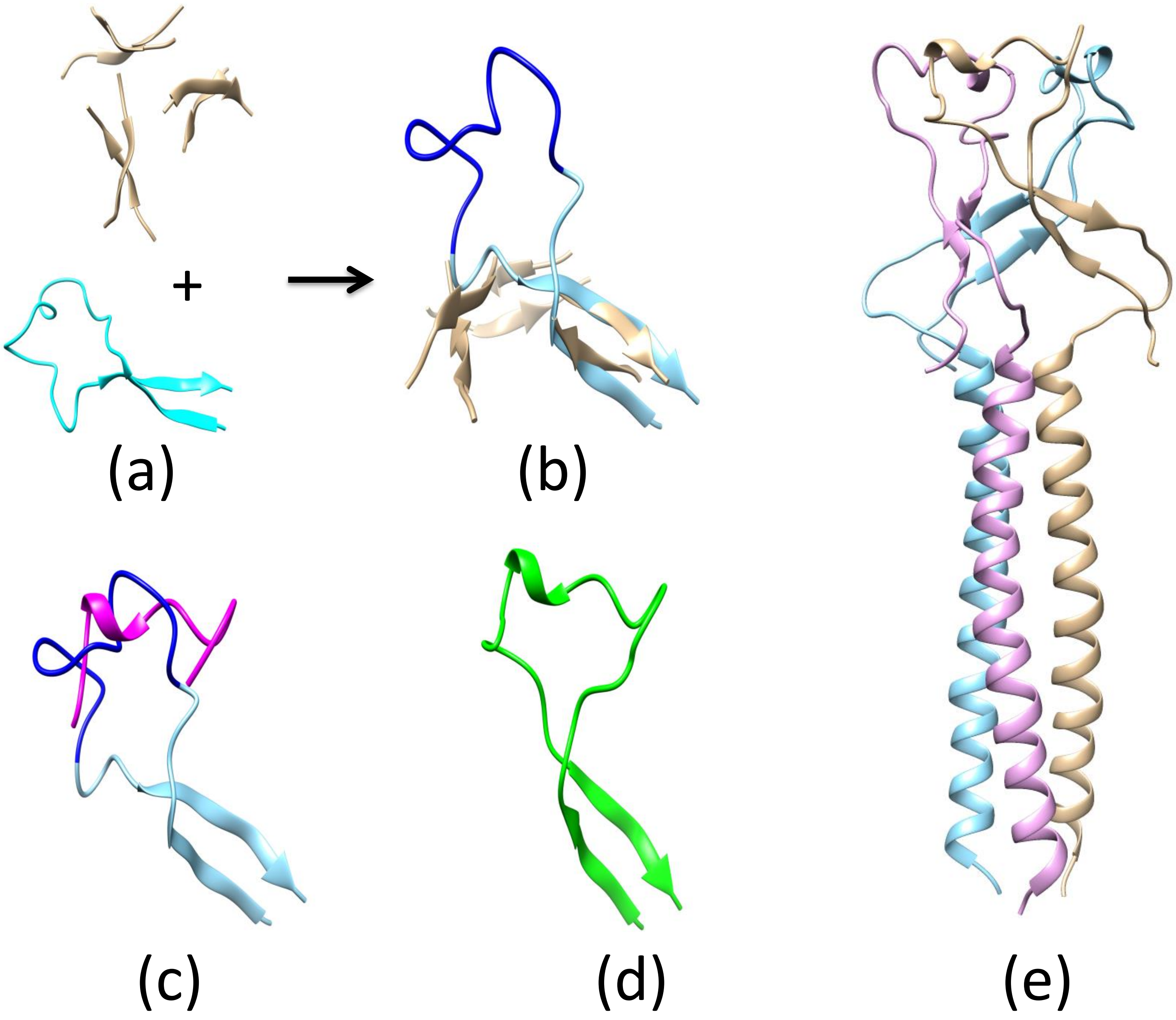

### Supplementary Materials

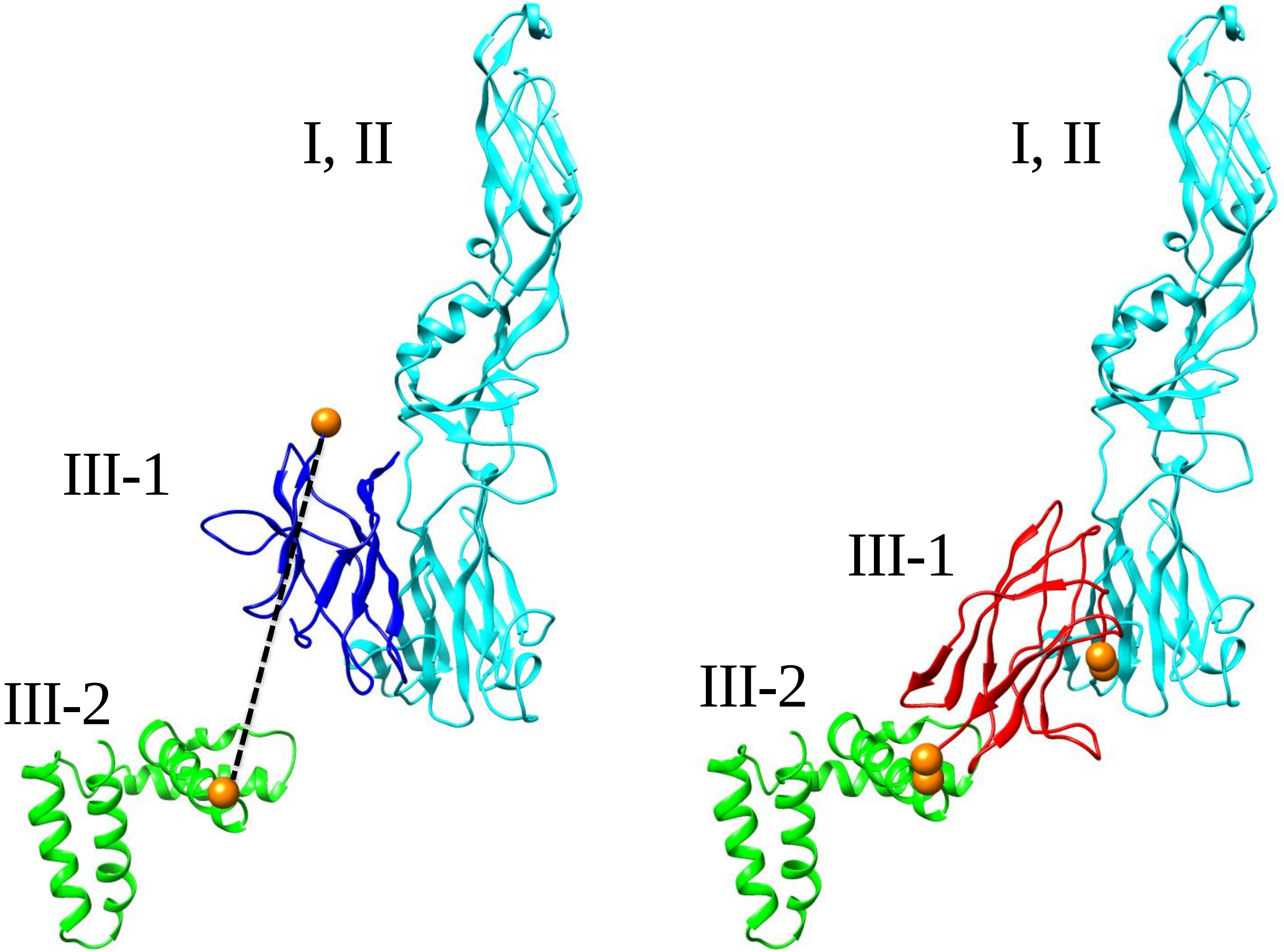


(a) (b)

### Supplementary Materials

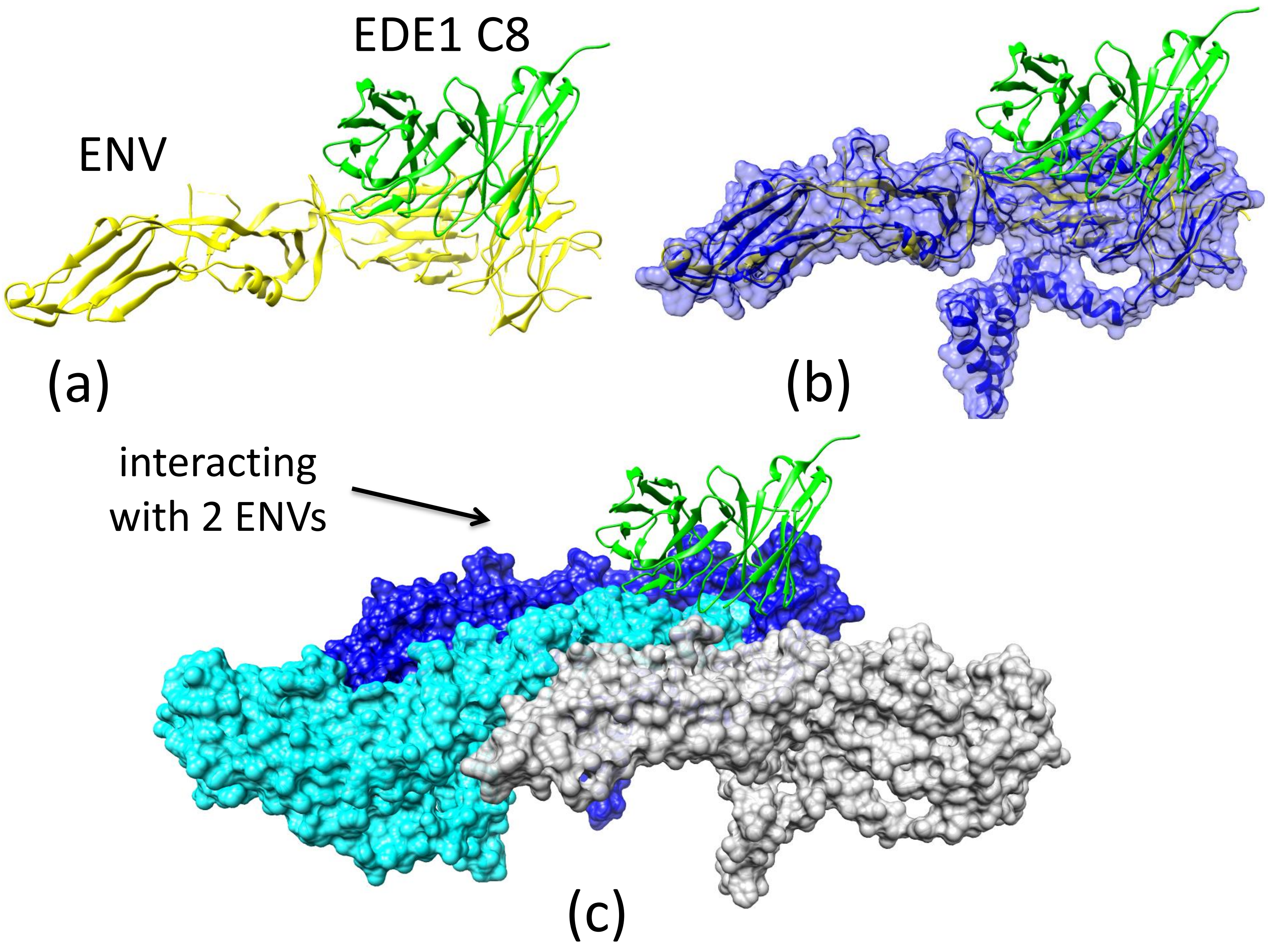
